# Cerebro: Interactive visualization of scRNA-seq data

**DOI:** 10.1101/631705

**Authors:** Roman Hillje, Pier Giuseppe Pelicci, Lucilla Luzi

**Author notes:** Corresponding Author: RH and PGP.

## Abstract

**Summary:** Despite the growing availability of sophisticated bioinformatic methods for the analysis of single-cell RNA-seq data, few tools exist that allow biologists without bioinformatic expertise to directly visualize and interact with their own data and results. Here, we present Cerebro (<u>ce</u>ll <u>re</u>port <u>bro</u>wser), a Shiny- and Electron-based standalone desktop application for macOS and Windows, which allows investigation and inspection of pre-processed single-cell transcriptomics data without requiring bioinformatic experience of the user.

Through an interactive and intuitive graphical interface, users can i) explore similarities and heterogeneity between samples and cells clusters in 2D or 3D projections such as t-SNE or UMAP, ii) display the expression level of single genes or genes sets of interest, iii) browse tables of most expressed genes and marker genes for each sample and cluster.

We provide a simple example to show how Cerebro can be used and which are its capabilities. Through a focus on flexibility and direct access to data and results, we think Cerebro offers a collaborative framework for bioinformaticians and experimental biologists which facilitates effective interaction to shorten the gap between analysis and interpretation of the data.

**Availability:** Cerebro and example data sets are available at https://github.com/romanhaa/Cerebro. Similarly, the R packages *cerebroApp* and *cerebroPrepare* R packages are available at https://github.com/romanhaa/cerebroApp and https://github.com/romanhaa/cerebroPrepare, respectively. All components are released under the MIT License.

## Introduction

Transcriptomics data of single cells (scRNA-seq) are generated with unprecedented frequency due to the recent availability of fully commercialized workflows and improvements in throughput and costs [1]. Though sophisticated bioinformatic tools are being developed [2, 3, 4], appropriate analyses and interpretation of data from scRNA-seq experiments largely relies on deep understanding of the biological context and experimental conditions behind the preparation of input cells. However, direct interaction with their own data sets is often out of reach for biologists without bioinformatic expertise. Cerebro aims to overcome the technical hurdles and allow direct and interactive exploration of pre-processed scRNA-seq results.

## Materials and methods

The key features of Cerebro include: i) visualization of two- and three-dimensional projections such as t-SNE or UMAP; ii) overview panels for samples and clusters; iii) tables of most expressed genes and marker genes for each sample and cluster; iv) tables of enriched pathways in marker genes of samples or clusters; v) visualization of expression of user-specified genes and gene sets from MSigDB [5, 6]. All these elements are designed to be interactive. Plots can be exported to PNG and/or PDF, while tables can be saved to CSV and Excel format.

The core of Cerebro is the *cerebroApp* application (built with Shiny [7]), which can be installed as a standalone application (built with Electron [8]). Alternatively, the Cerebro user interface is available as an R package or a Docker container. Input data needs to be prepared using the *cerebroPrepare* R package, which is separated from the *cerebroApp* package to minimize software dependencies. Currently, the *cerebroPrepare* package offers functionality to export a Seurat object (both Seurat v2 and v3 are supported) to the correct format in a single step [9]. Furthermore, *cerebroPrepare* provides functions to perform a set of (optional) analyses, e.g. pathway enrichment analyses based on marker gene lists of samples or clusters through Enrichr [10, 11]. Parallel processing in these functions ensures time-efficient execution. For human and mouse data sets, marker genes will be intersected with the gene ontology term “cell surface” (GO:0009986) to highlight potential markers for enrichment of the respective cell community. The exported .*crb* file is then loaded into Cerebro and shows the information from the Seurat object (Figure 1). Full-size versions of the examples of the Cerebro interface shown in Figure 1 can be found in the supplementary figures as well as in the Cerebro GitHub repository.

**Figure 1:**
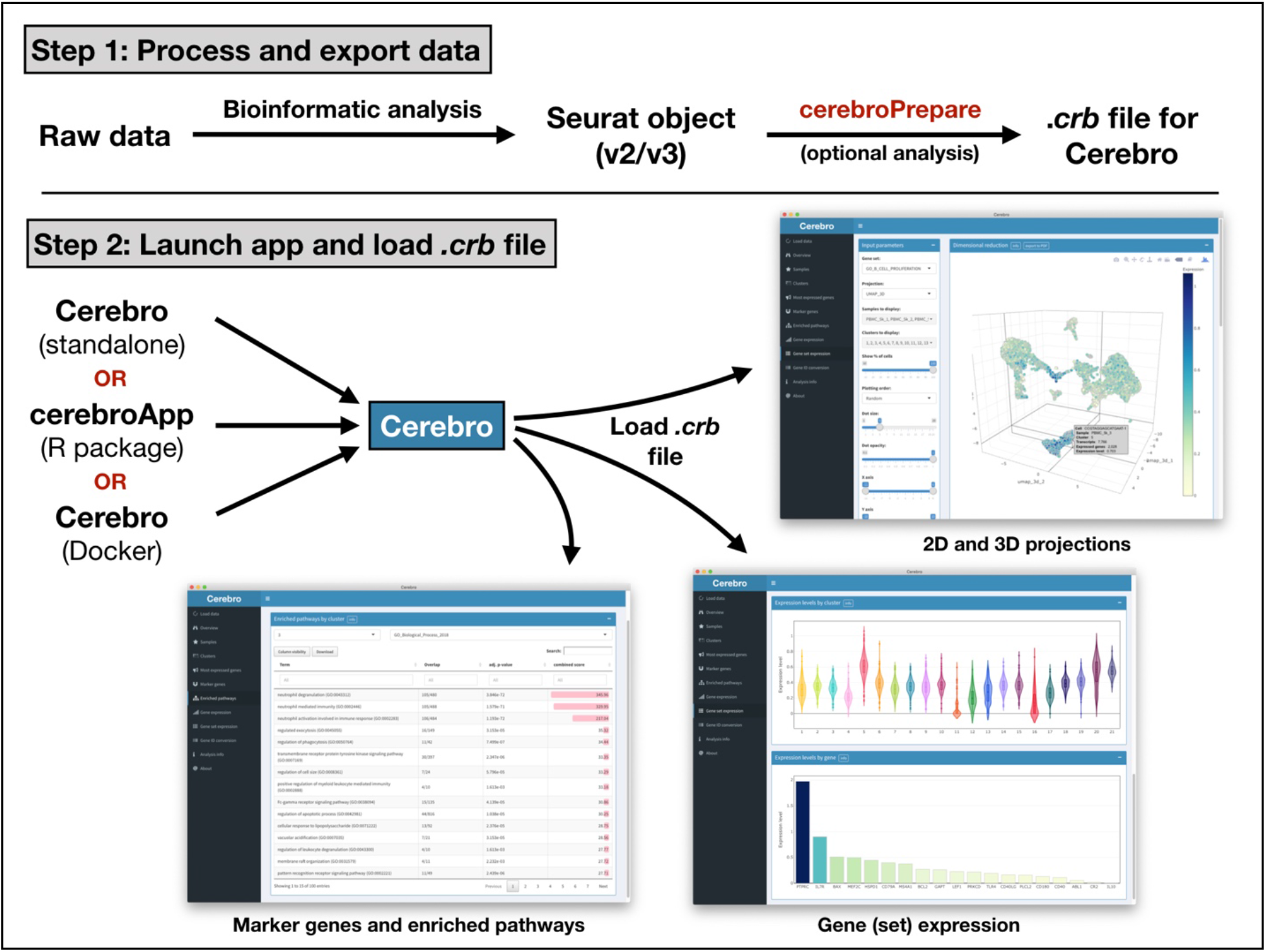
Schematic workflow of Cerebro. In the first step, the raw data is processed and analyzed (barcode extraction, alignment, etc.), stored in a Seurat object and exported to a .*crb* file using the *cerebroPrepare* package. Subsequently, the .*crb* file can then be loaded into Cerebro for visualization. Currently, Cerebro can be launched as a standalone application, as the corresponding *cerebroApp* R package or from the dedicated Docker container.

## Usage scenario

To illustrate the proposed workflow, we analyzed a publicly available scRNA-seq data set containing ~10k human PBMCs from a healthy donor (link to data set in Supplementary Information) following the basic Seurat workflow. First, we loaded the feature matrix and created a Seurat object, filtered cells based on numbers of transcripts and expressed genes, log-transformed transcript counts and normalized each cell to 10,000 transcripts. We then identified variable genes, scaled the expression matrix and regressed out numbers of transcripts, performed cell cycle and principal component analysis, identified clusters and described their relationship in a cluster tree. We also generated 2D and 3D projections using the tSNE and UMAP algorithms. Then, we used *cerebroPrepare* to calculate the percentage of mitochondrial and ribosomal gene expression, obtain the most expressed genes and marker genes for each sample and cluster, perform pathway enrichment analysis using the identified marker genes, and finally export a *.crb* file that can be loaded into Cerebro.

Based on the combined information from pathway enrichment, marker genes and expression of additional genes and gene sets, we were able to retrieve expected cell types commonly found in PBMC samples (dendritic cells, NK cells, B cells, megakaryocytes, monocytes, CD4+ and CD8+ T cells) and assign a cell type to each cluster. If desired, these cell groups could be further discriminated by checking the expression of additional marker genes and gene sets. To give a better example of the functionality of Cerebro, we randomly split the example data set, which consists of cells from a single sample, into three samples to simulate a dataset with multiple samples.

## Conclusion

By providing access to comprehensive information on expression profiles of samples and clusters, we hope that Cerebro will accelerate data interpretation and ultimately knowledge acquisition. Notably, the proposed workflow also provides analytical flexibility by enabling the addition of custom analyses and results to the Seurat object. Since the code is completely open-source, it is possible (and people are encouraged) to modify and adapt Cerebro to display other results and data types. While *cerebroPrepare* currently only supports to prepare Seurat objects for visualization in Cerebro, export methods for object types of other popular scRNA-seq analysis frameworks, such as *SingleCellExperiment* or *AnnData* (used by scanpy [12]) can be added in the future. Furthermore, Seurat already provides functionality to import data from other frameworks, including the two mentioned above, and therefore serves as a gateway for the majority of data sets. Due to the nature of Shiny apps, Cerebro can be easily adapted to be hosted on web servers.

## Software availability

The current Cerebro release for Windows and macOS, together with the source code, can be downloaded from the GitHub repository: https://github.com/romanhaa/Cerebro/releases. Source code of the *cerebroApp* R package and Shiny application on which Cerebro is based can be accessed at and installed from: https://github.com/romanhaa/cerebroApp. Source code of the *cerebroPrepare* R package containing functions for data preparation and processing can be found at: https://github.com/romanhaa/cerebroPrepare. Analysis of the example data set was carried out in a Docker container to ensure reproducibility. The container is available through the Docker Hub and was built using a recipe file stored in the Cerebro GitHub repository: https://cloud.docker.com/u/romanhaa/repository/docker/romanhaa/cerebro-example

## Acknowledgements

We thank our colleagues and early version users for their constructive feedback.

## Author contributions

RH: Conceptualization; Development; Writing - Original Draft

LL: Conceptualization; Supervision; Writing - Original Draft

PGP: Supervision; Writing - Original Draft; Funding Acquisition

## Supplementary information

The example data set was generated using the 10x Genomics Chromium Single Cell 3’ Library & Gel Bead Kit v3 and processed using Cell Ranger 3.0.0. It is available through the 10x Genomics website (https://support.10xgenomics.com/single-cell-gene-expression/datasets/3.0.0/pbmc_10k_v3) and licensed under the Creative Commons Attribution 4.0 International license (https://creativecommons.org/licenses/by/4.0/).

The .*crb* files from the example data set (Seurat v2 and v3), as well as the scripts and raw data can be found in the Cerebro GitHub repository (https://github.com/romanhaa/Cerebro). The entire workflow described above took roughly 25 minutes to finish on our HPC.

Further screenshots and explanation are available in the Cerebro GitHub repository as well (https://github.com/romanhaa/Cerebro).

**Supplementary figure 1:**
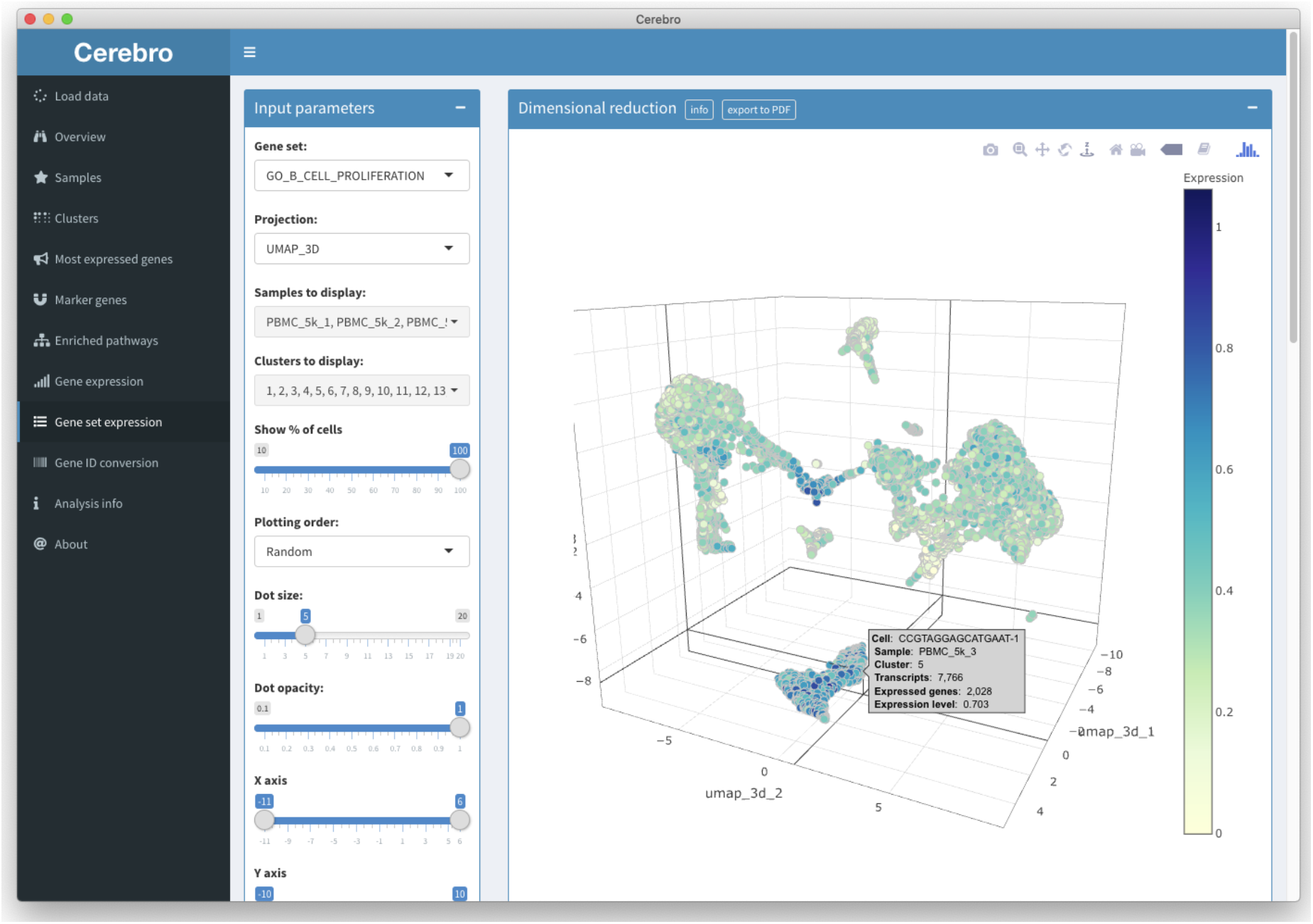
Gene set expression in 3D projection. The “Gene set expression” panel gives access to the gene sets available in the Molecular Signatures Database (MSigDB). Shown here is the average expression of the genes of the “GO_B_CELL_PROLIFERATION” signature in the 3D UMAP projection. The plot can be rotated, and each cell can be identified by the cursor, showing additional information such as its name/barcode and the sample and cluster it belongs to.

**Supplementary figure 2:**
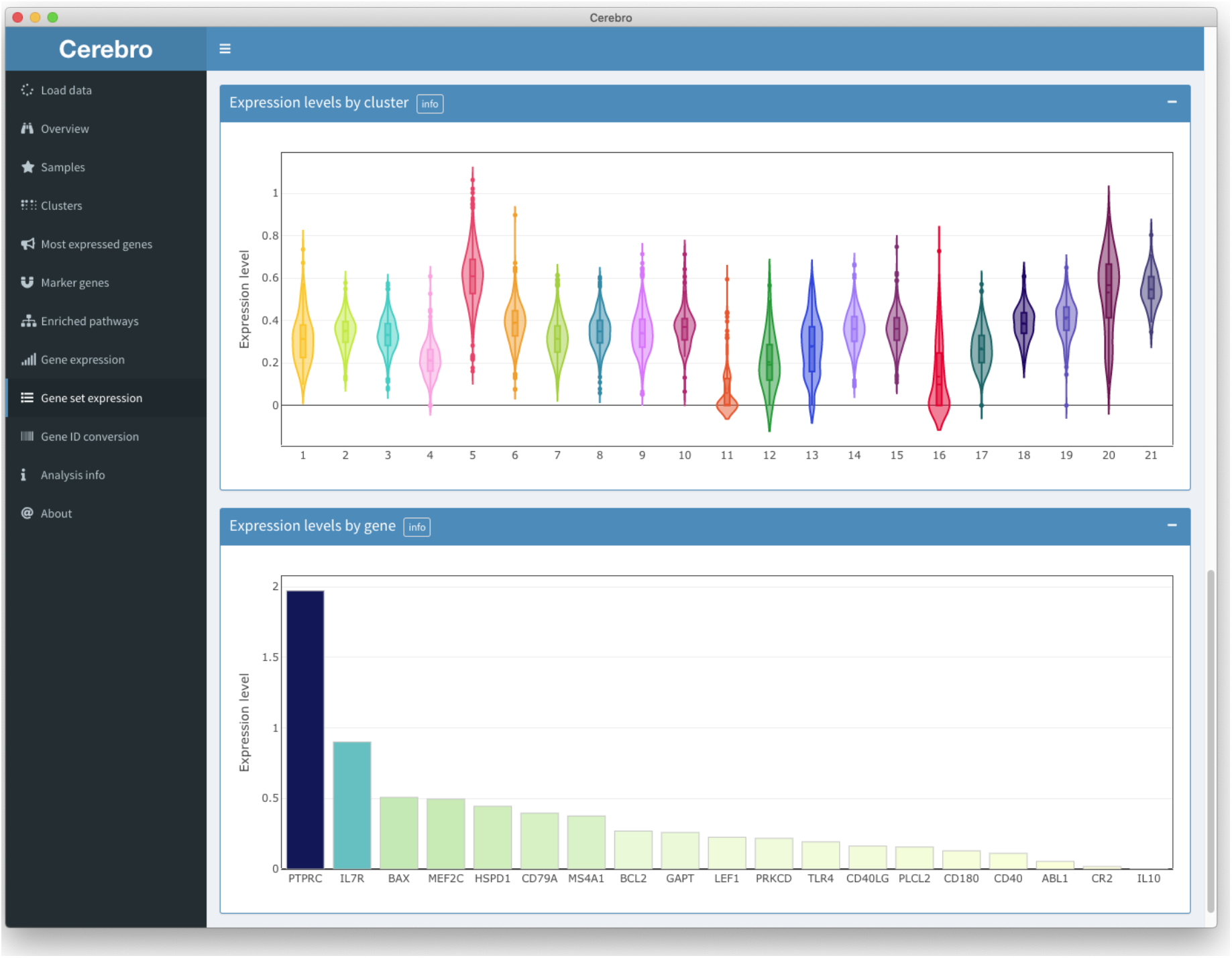
Gene set expression by cluster. The “Gene set expression” panel gives access to the gene sets available in the Molecular Signatures Database (MSigDB). Shown here is the average expression of the genes of the “GO_B_CELL_PROLIFERATION” signature for every cluster, as well as the average expression of each individual gene of the selected set across all cells, ranked from high to low. This allows to identify genes that potentially overshadow the contribution of other genes due to overall stronger expression.

**Supplementary figure 3:**
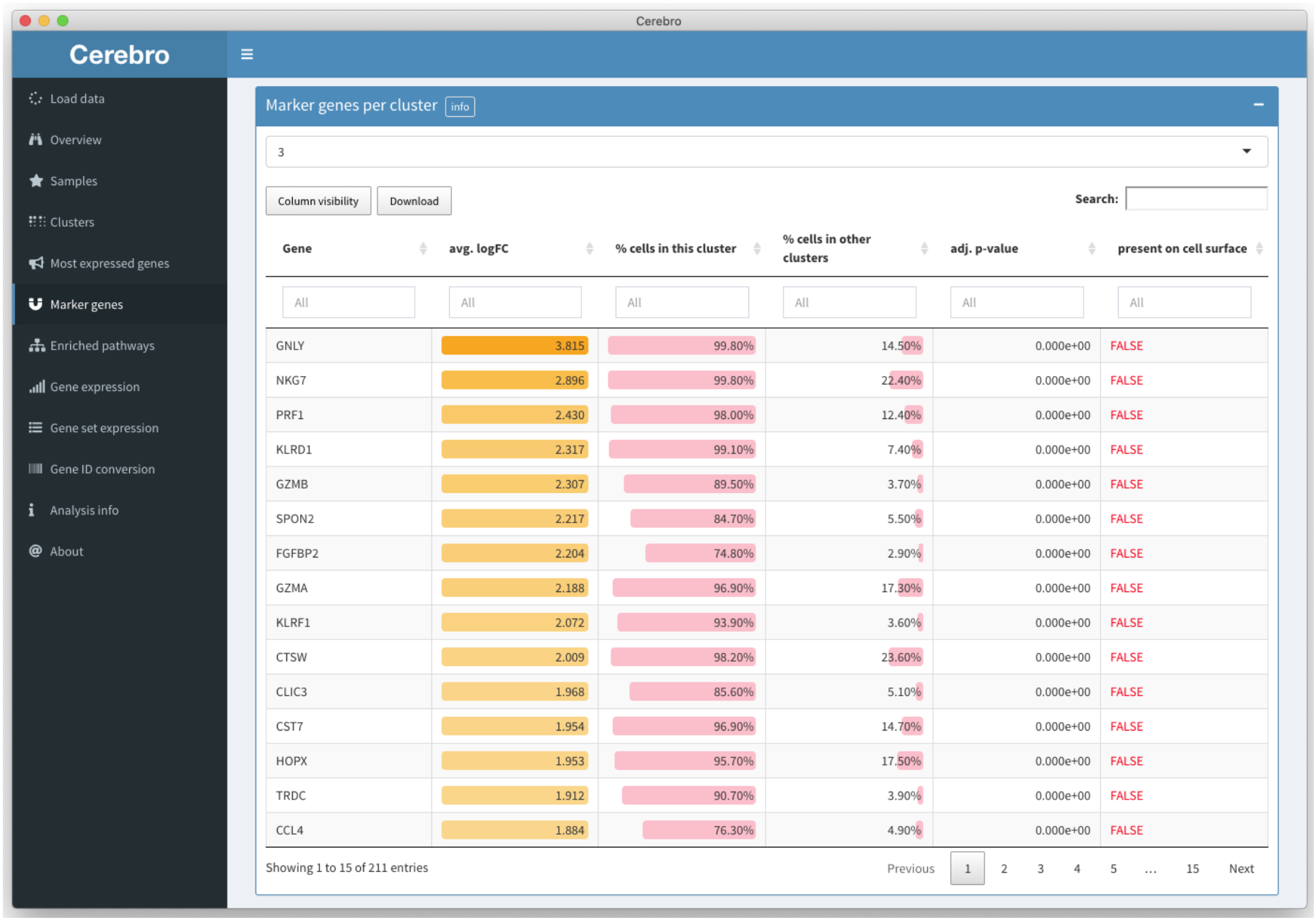
Table of marker genes. The “Marker genes” panel shows tables of marker genes for each sample and cluster. Shown here are the marker genes for cluster 5. The output can be filtered and sorted by the average log-fold change, the percentage of cells in the selected cluster that express the gene, the percentage of cells in the rest of the cells that express the gene, and the adjusted p-value. The last column indicates if a gene is part of the gene ontology term “cell surface” (GO:0009986) to facilitate the identification of cell surface markers for enrichment of the selected cell community.

**Supplementary figure 4:**
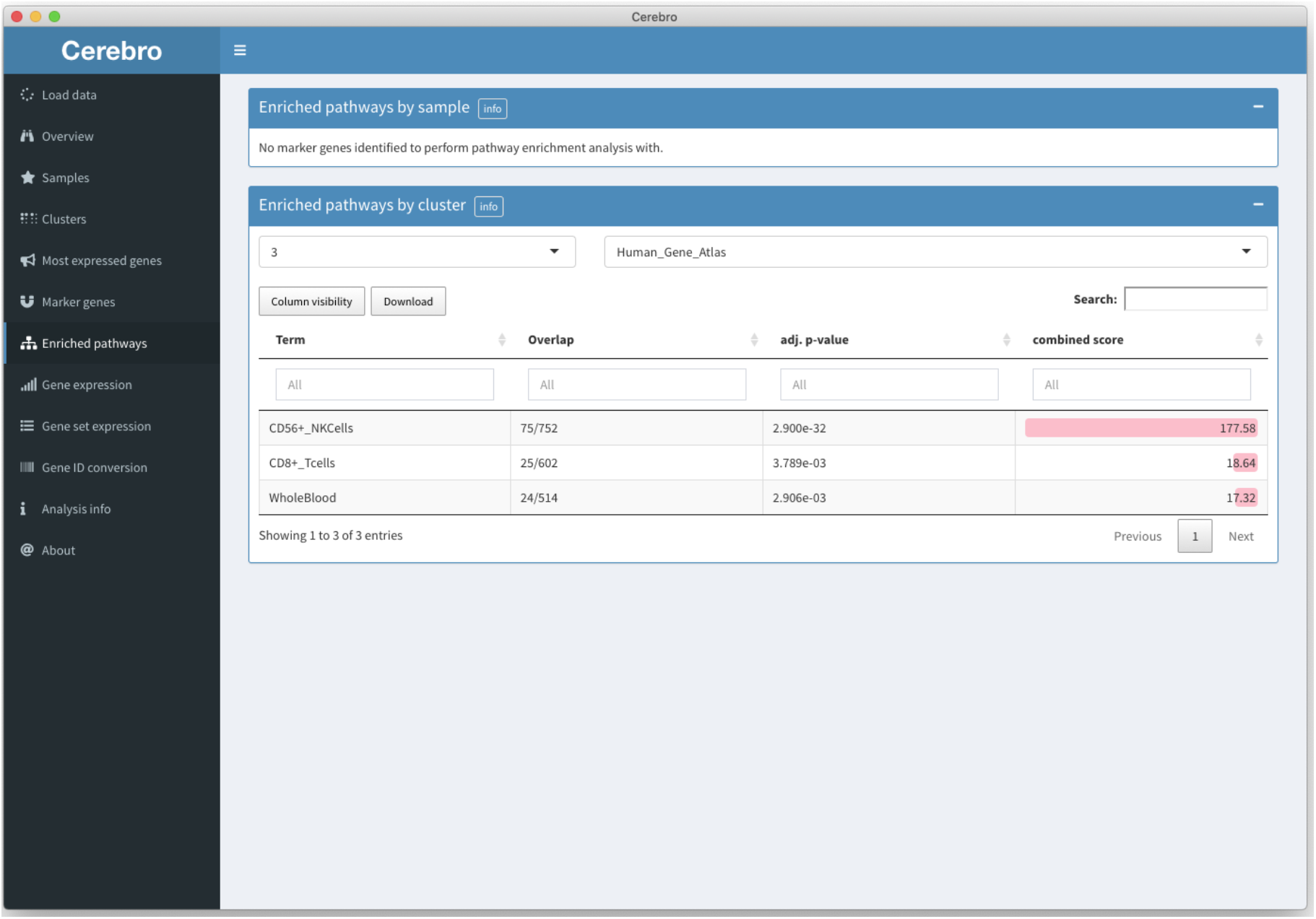
Table of enriched pathways in marker genes. The “Enriched pathways” panel shows the results of the pathway enrichment analysis based on marker genes of each sample and cluster. Through the Enrichr web service, multiple databases are queried. Shown here are the enrichment results from the “Human_Gene_Atlas” database for cluster 3. As indicated by the combined score, the cells of this cluster are likely NK cells. The genes that overlap with each term can be listed in an additional column that is hidden by default to improve visibility.

